# Infection drives localised and individualised antibody repertoires, whereas immunisation promotes repertoire convergence in pigs

**DOI:** 10.64898/2026.05.15.725346

**Authors:** Emily J Briggs, Shafiyeel Chowdhury, Bharti Mittal, Valentine Chaillot, Charlotte May, Kevan Hanson, Ehsan Sedaghat-Rostami, Sanis Wongborphid, Erica Bickerton, Basudev Paudyal, Christine S. Rollier, Cornelis A.M. de Haan, Kristien Van Reeth, Sarah Keep, Matthew D. J. Dicks, Danish Munir, Marie Bonnet-Di Placido, Elma Tchilian

## Abstract

Respiratory viruses elicit mucosal and systemic immunity, yet how tissue compartmentalisation and exposure route shape B cell repertoires remain unclear. Using the pig model, we characterised antibody repertoires in bronchoalveolar lavage, tracheobronchial lymph nodes, spleen, and blood following intranasal infection with pandemic influenza virus (pH1N1) or porcine respiratory coronavirus (PRCV), and after intramuscular PRCV immunisation. Infection induced highly compartmentalised responses, with antigen-associated clones enriched in lung, lymph nodes and spleen but with limited presentation in blood. These clones were largely private, indicating individualised responses, and displayed tissue-specific diversity. The similar tissue distributions between pH1N1 and PRCV infections suggest a conserved spatial organisation, although differential lymph node involvement indicates pathogen-specific effects. In contrast, intramuscular immunisation generated more convergent, public repertoires in blood, characterised by reduced clonal size diversity. Monoclonal antibody analysis revealed functional heterogeneity and limited overlap with bulk repertoires. Together, these findings show that despite stable germline usage, functional antibody responses are shaped by the route of antigen exposure, clonal selection, and host specific factors, resulting in distinct repertoire architectures between infection and immunisation.

**Author summary:** Respiratory viruses such as influenza and coronaviruses infect the airways and lungs, where they trigger immune responses that help the body fight infection. Most studies of immunity rely on blood, although many immune cells function in tissues like the lungs. Therefore, we may be missing important parts of the immune response.

We used pigs, a large animal model that closely resembles humans in respiratory biology and is naturally infected with influenzas and coronaviruses, to investigate how immune responses are shaped by infection compared with intramuscular vaccination. We examined B cells in lungs, nearby lymph nodes, spleen, and blood after infection with influenza or a porcine coronavirus and compared these with responses after vaccination.

We found that infection generates distinct, individual immune responses in the lung and lymphoid organs. Vaccination by injection led to more similar responses across animals, mostly in the blood. This suggests the way the immune system encounters an antigen influences the response generated.

We also showed that antibodies with similar binding properties can behave very differently in their ability to neutralise viruses. Our findings show that infection and vaccination shape immunity in fundamentally different ways, with important implications for designing vaccines that better protect against respiratory diseases.

## Introduction

Respiratory viral infections remain a major global health burden, with influenza A viruses (IAV) and coronaviruses (CoV) causing periodic epidemics and pandemics. Understanding how these viruses shape the B cell repertoire across tissues is important, as they engage both mucosal and systemic immunity. Circulating antibodies are widely used as a proxy for immune status, yet they represent only a fraction of the total response, with a substantial component residing in tissues such as the respiratory tract and its draining lymph nodes. Much less is known about the repertoire of tissue resident B cells, particularly in the lung^1^. Studies of influenza and COVID-19 have begun to address this gap, showing that lung resident memory B cells are established locally following infection and exhibit distinct transcriptional and phenotypic profiles compared with circulating populations^2^. These cells are retained within the lung and airways, do not recirculate, and can rapidly differentiate into antibody secreting cells upon re-exposure, supporting local antibody production at sites of infection^3^. Together, these findings suggest that mucosal antibody repertoires may be distinct from those in peripheral blood, either due to local antigen driven selection or differences in B cell composition and maturation within tissues. However, the extent to which clonal expansion, repertoire diversity, and inter individual sharing are shaped by tissue and route of antigen exposure remains unclear. Defining the organisation of compartmentalised responses, and how they differ following infection with IAV or CoV or parenteral immunisation, is essential for understanding protective immunity and guiding vaccine design^4^.

The domestic pig is a highly relevant large natural host animal model for studying respiratory virus infections, particularly IAV and CoV^5-8^. Pigs share many anatomical, physiological, and immunological features with humans, including similar lung structure, airway architecture, lung surface area, and distribution of airway immune cells^9-11^. They also exhibit comparable distribution of sialic acid receptors, which contributes to their natural susceptibility to IAV that are antigenically and genetically related to human strains, resulting in clinical signs, lung pathology, and immune responses similar to those observed in humans^8, 12, 13^. Pigs are natural hosts for several CoV, including porcine respiratory coronavirus (PRCV), which induces lung pathology comparable to that in patients with severe acute respiratory syndrome (SARS)-CoV and SARS-CoV-2^5, 14^. These similarities, together with their role as natural hosts of IAV and CoV with zoonotic potential, make pigs an excellent translational model for understanding host pathogen interactions and evaluating novel vaccines and therapeutics.

The immunoglobulin heavy chain locus in pigs is characterised by relatively restricted germline diversity, with a limited number of functional IGHV segments^15^. Classical work by John E. Butler and more recent studies by Driver et al. demonstrate that only a small subset of V genes contributes to most of the antibody diversity^1, 16, 17^. Because of the small number of IGHV, IGHD, and IGHJ segments used, the combinatorial diversity in pigs is comprised of only ∼14 possibilities compared to ∼9,000 in humans^1^. Consequently, repertoire diversification relies primarily on junctional diversity and somatic hypermutation rather than extensive germline gene usage. These features make the pig a useful comparative model for dissecting how antigen-driven selection and tissue specific factors shape antibody repertoire architecture in the context of limited germline variation.

In this study, using B cell sequencing in bronchoalveolar lavage, tracheobronchial lymph nodes, spleen and peripheral blood, we investigated the antibody repertoires in pigs following respiratory infection with either influenza virus or PRCV or following intramuscular PRCV immunisation. In addition, PRCV-specific monoclonal antibodies (mAbs) were generated to further assess the functional properties of the response. By integrating tissue distribution, inter-animal sharing, repertoire diversity, and monoclonal antibody characterisation, we show that respiratory infection is associated with localised and individualised repertoire responses, whereas intramuscular immunisation promotes a more convergent response across animals.

## Results

### Antibody repertoires following pH1N1 or PRCV Blg20 infection

Ten pigs each were infected with either A/swine/Gent/53/2019 (pH1N1) or PRCV/swine/Belgium/PS-071/2020 (PRCV Blg20). At 5 and 12 days post-infection, 5 animals per infection group and two control uninfected animals were culled. IgG and IgA sequences from bronchoalveolar lavage (BAL), tracheobronchial lymph node (TBLN), peripheral blood mononuclear cells (PBMC) and spleen samples from animals culled 5 and 12 days post-infection were sequenced for antibody repertoire analysis (**Supp. Fig. S1A**). Samples underwent quality control (**Supp. Fig. S1B, C**) and reads were normalised to enable equal representation of clonotypes across samples with different sequencing depths.

Clusters were defined as groups of heavy chain sequences sharing at least 94% identity across the full variable region, representing closely related antibody sequences. Clusters were classified as ‘selected’ (potentially antigen-specific) or ‘irrelevant’ (unlikely to be antigen-specific) based on their kinetics: clusters were classified as irrelevant if at least 10% of normalised reads were derived from pre-bleed samples or uninfected control animals. All other clusters were classified as selected (**Supp. Fig. S1D, E**). These selected clusters should therefore be interpreted as kinetically enriched, response-associated repertoire features rather than as definitively antigen-specific clones.

### Tissue distribution, publicity, and diversity of clusters following pH1N1 infection

Following pH1N1 infection, irrelevant clusters contained 2.1-fold more sequences from PBMC (51.1%) compared to selected clusters (24.3%) (**Fig. 1A**). In contrast, selected clusters were enriched in BAL, TBLN or spleen between 1.8- and 2.4-fold compared to irrelevant clusters (Χ^2^ = 14912, df = 4, p < 0.0001) (**Fig. 1B, Supp. Table S1**). Clusters containing sequences from two or more tissues were defined as shared clusters. The overall proportion of shared clusters was similar between groups (26% of irrelevant and 27% of selected clusters), although distributed differently between tissues. These results showed that clusters associated with response to pH1N1 infection were preferentially localised to BAL, TBLN and spleen rather than the peripheral blood.

**Figure 1:**
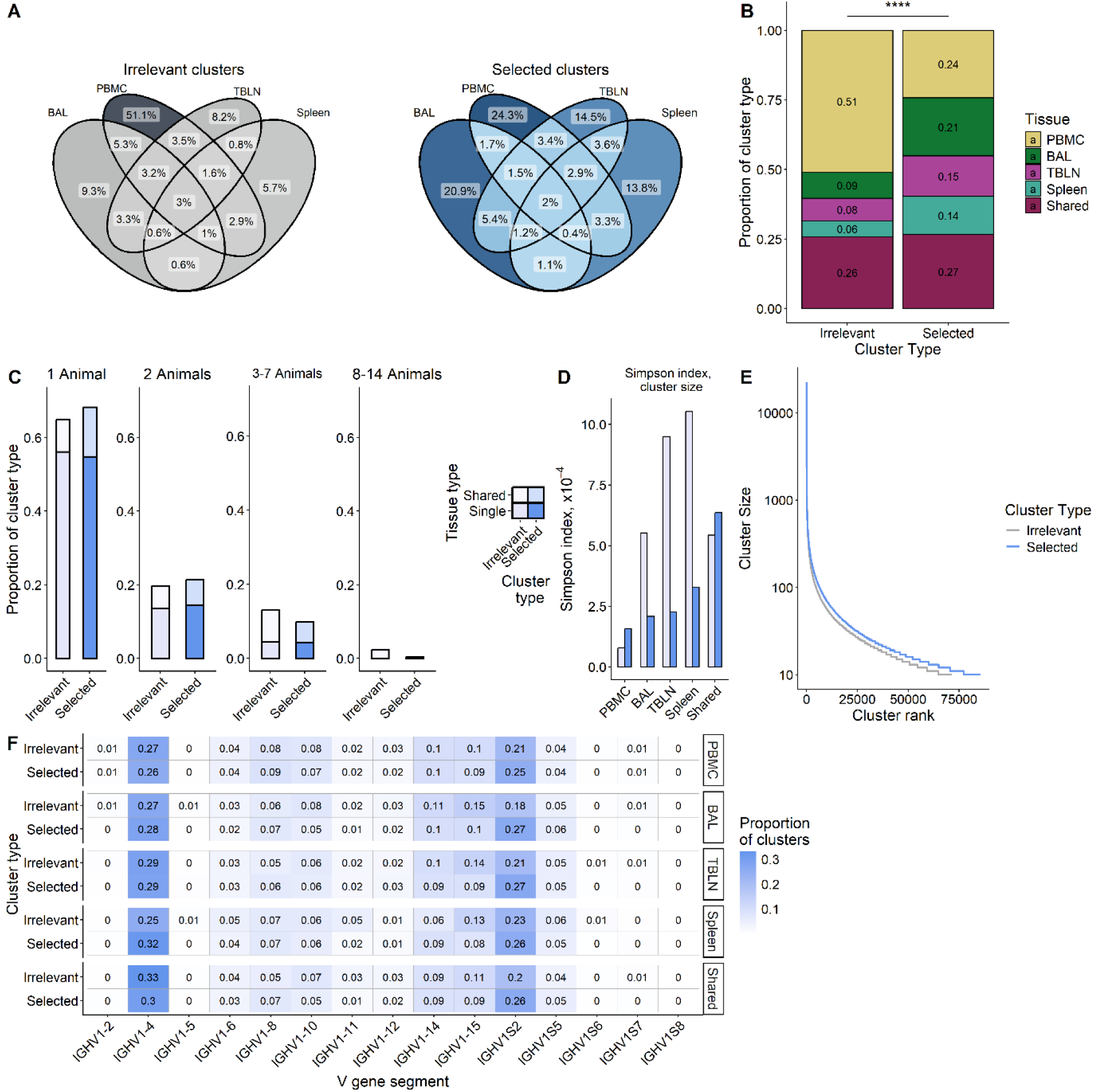
Antibody repertoire following infection with pH1N1 Selected and irrelevant clusters were analysed for their tissue distribution, sharing between animals, diversity, size, and V gene segment usage following pH1N1 infection. **A:** Distribution of irrelevant (grey) and selected (blue) clusters across tissues. Percentages of clusters of each type are indicated. **B:** Proportion of irrelevant and selected clusters containing sequences from a single tissue, or from more than one tissue. Distribution across tissues was analysed by Chi-square test (Χ2 = 13206, df = 4, p < 2.2 x10-16). **C:** Publicity of irrelevant and selected clusters. Clusters were grouped according to whether they contained sequences from one animal, two animals, 3-7 animals, or 8-14 animals and were further classified as restricted to a single tissue (Single) or detected across two or more tissues (Shared). **D:** Diversity of cluster size in irrelevant and selected clusters was determined by Simpson index. Alower Simpson index indicates greater diversity. **E:** Size distribution of irrelevant and selected clusters. **F:** V gene segment usage in centroid sequences of irrelevant and selected clusters that were either restricted to single tissues (PBMC, BAL, TBLN, or spleen) or shared across two or more tissues. **** indicates p < 0.0001.

We next assessed the extent to which clusters were shared between individuals (publicity of clusters). Overall, irrelevant clusters were more public than selected clusters, particularly among shared clusters (Χ^2^ = 2303.2, df = 3, p < 0.0001). Shared irrelevant clusters more frequently contained sequences from multiple animals, with 0.26% detected in 14 animals. By contrast, only 0.03% of selected clusters were detected across multiple tissues and all 14 animals (X^2^ = 162.98, df = 1, p < 0.001). Selected clusters were more often private or shared between only two animals, particularly amongst clusters detected in only one tissue (Χ^2^ = 40.143, df = 3, p < 0.0001) (**Fig. 1C, Supp. Table S2, S3**). This suggest that multi-tissue irrelevant clusters may reflect widely shared background repertoire, whereas multi-tissue dissemination of selected clusters was largely animal-specific, indicating individualised responses to pH1N1 infection.

Cluster size diversity was quantified using Simpson index. Selected clusters restricted to BAL, TBLN, or spleen showed greater diversity in cluster size than irrelevant clusters, indicated by decreased Simpson index (**Fig 1D**). PBMC-only and shared selected clusters showed reduced size diversity compared to irrelevant clusters (**Fig. 1D**). Furthermore, selected clusters were generally larger than irrelevant clusters (**Fig. 1E**). Positional diversity was analysed using the centroid amino acid sequence of each cluster, defined as the heavy chain sequence with the greatest connectivity to other sequences within that cluster. Positional diversity of selected and irrelevant clusters was assessed using Shannon index. Selected clusters in the PBMC and spleen exhibited increased positional diversity compared to irrelevant clusters, with a modest increase also observed in TBLN. A modest decrease in positional diversity was observed in selected clusters restricted to the BAL and those shared across tissues (**Supp. Fig. S1F**). Together, these data indicate that pattern of diversity in selected clusters is not uniform across tissues, instead showing tissue-specific differences in both cluster size distribution and amino acid sequence diversity.

V gene segment usage was analysed separately in selected and irrelevant clusters that were either restricted to a single tissue or shared across multiple tissues. Two V genes, IGHV1-4 and IGHV1S2, comprised approximately 50% of both selected and irrelevant clusters across all tissues. The next most dominant V gene segments were IGHV1-15, IGHV1-14, and IGHV1-8. Together, these five gene segments comprised approximately 75% of the repertoire (**Fig. 1F**). No treatment- or tissue-specific differences in V gene usage were observed, consistent with previous reports^18^.

Overall, following pH1N1 infection, antigen-associated selected clusters were enriched in BAL, TBLN and spleen, with reduced representation in peripheral blood, indicating tissue-localised responses. These clusters were more private and exhibited tissue-specific patterns in cluster size and sequence diversity, while V gene usage was dominated by a limited set of genes.

### Tissue distribution, publicity, and diversity of clusters following PRCV Blg20 infection

Cluster distribution across tissues was analysed following PRCV Blg20 infection. Irrelevant clusters were more frequently restricted to PBMC than selected clusters (1.9-fold). Clusters restricted to BAL and spleen were enriched 2.6- and 2.4-fold respectively in selected compared to irrelevant clusters (Χ^2^ = 13206, df = 4, p < 0.0001). However, no enrichment of TBLN in selected clusters was observed (**Fig. 2A**). Proportion of clusters detected across multiple tissues was similar between groups, (23% of irrelevant clusters and 25% of selected clusters) (**Fig. 2B, Supp. Table S4**). Overall, PRCV Blg20 infection was associated with reduced representation of clusters in the PBMCs and enrichment of selected clusters in BAL and spleen, but not in TBLN.

**Figure 2:**
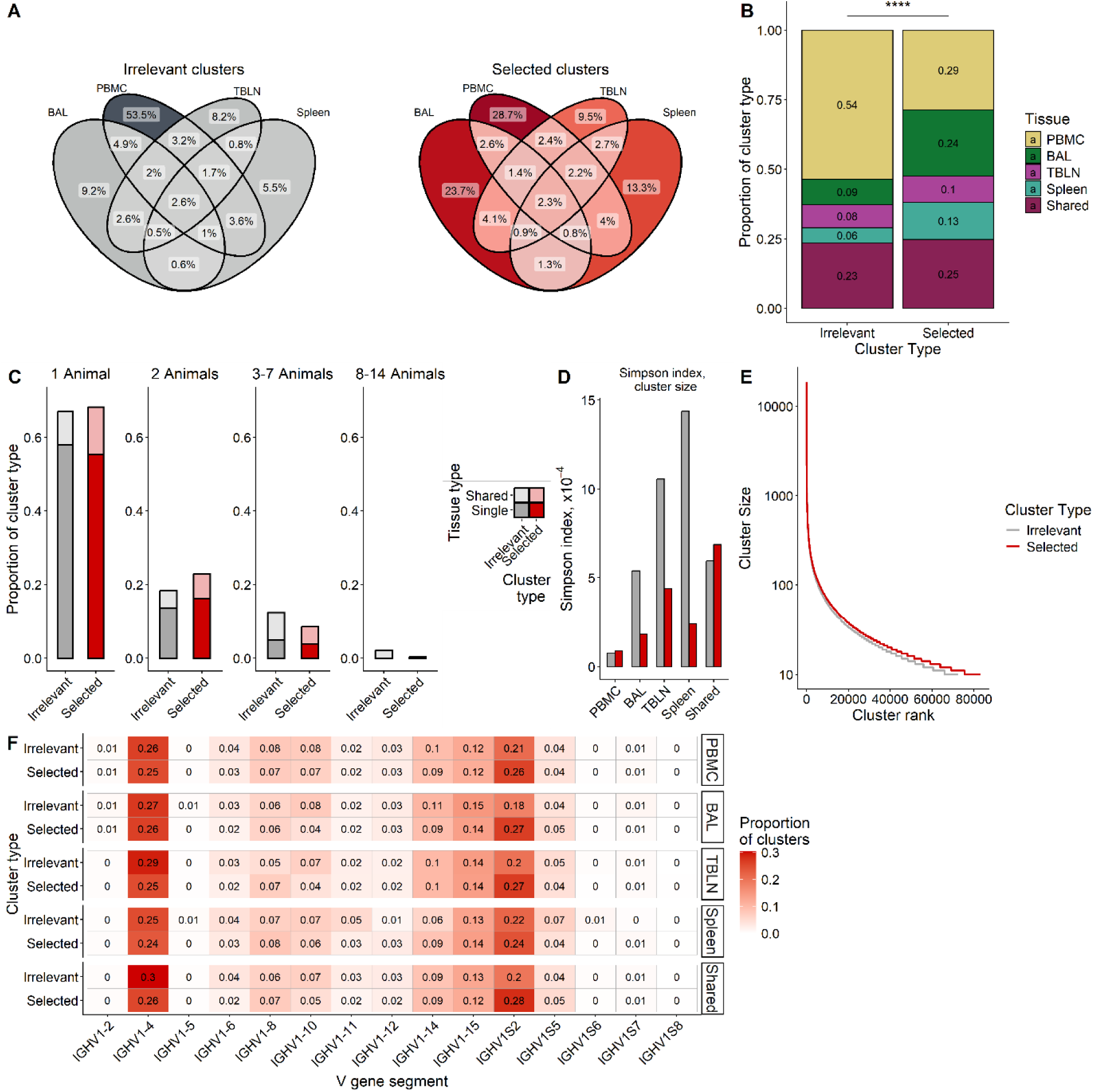
Antibody repertoire following infection with PRCV Blg20 Selected and irrelevant clusters were analysed for their tissue distribution, sharing between animals, diversity, size, and V gene segment usage following PRCV Blg20 infection. **A:** Distribution of irrelevant (grey) and selected (blue) clusters across tissues. Percentages of clusters of each type are indicated. **B:** Proportion of irrelevant and selected clusters containing sequences from a single tissue, or from more than one tissue. Distribution across tissues was analysed by Chi-square test (Χ2 = 13206, df = 4, p < 2.2 x10-16). **C:** Publicity of irrelevant and selected clusters. Clusters were grouped according to whether they contained sequences from one animal, two animals, 3-7 animals, or 8-14 animals and were further classified as restricted to a single tissue (Single) or detected across two or more tissues (Shared). **D:** Diversity of cluster size in irrelevant and selected clusters was determined by Simpson index. Alower Simpson index indicates greater diversity. **E:** Size distribution of irrelevant and selected clusters. **F:** V gene segment usage in centroid sequences of irrelevant and selected clusters that were either restricted to single tissues (PBMC, BAL, TBLN, or spleen) or shared across two or more tissues. **** indicates p < 0.0001.

Distribution of irrelevant cluster publicity was analysed in the PRCV Blg20 repertoire. A small fraction of irrelevant clusters (190, 0.26%) was highly public, containing sequences from multiple tissues and all 14 animals compared to 10 in selected clusters (0.012%) (Χ^2^ = 188.16, df = 1, p < 0.0001). In contrast, selected clusters shared across tissues were more often restricted to a single animal than irrelevant clusters (Χ^2^ = 2190.8, df = 3, p < 0.0001) (**Fig. 2C, Supp. Table S5, S6**).

Selected clusters restricted to BAL, TBLN or spleen exhibited higher diversity in cluster size than irrelevant clusters, indicated by decreased Simpson index (**Fig. 2D**). Positional amino acid diversity was increased in selected clusters from PBMC, BAL, and TBLN, with little difference observed in spleen-only or shared clusters (**Supp. Fig. 1I**). Cluster size was generally larger in the selected clusters compared to irrelevant (**Fig. 2E**). Together, these findings indicate that selected clusters in the PRCV Blg20 repertoire were associated with increased diversity in both cluster size and amino acid sequence. This is consistent with a broader and more heterogenous repertoire associated with PRCV Blg20 infection compared to irrelevant clusters. V gene usage in the PRCV Blg20 repertoire was dominated by five V gene segments, with IGHV1-4, IGHV1S2, IGHV1-15, IGHV1-14, and IGHV1-8 comprising approximately 75% of the repertoire. Again, no tissue- or treatment-specific differences were observed (**Fig. 2F**).

Together, these findings indicate that selected clusters differed consistently from irrelevant clusters following PRCV Blg20 infection. Selected clusters were detected less frequently in PBMCs and more frequently in BAL and spleen, but not TBLN. Selected clusters were largely private and showed increased clonal and sequence diversity, while overall V gene usage remained restricted.

### Comparison of antibody repertoires following pH1N1 and PRCV Blg20 infection

We next compared the repertoires following pH1N1 and PRCV Blg20 infections. Tissue distribution patterns were broadly similar between the two infections (**Fig. 3A**). In both datasets, selected clusters were more frequently restricted to BAL and spleen and less frequently to PBMC compared to irrelevant clusters, while proportion of shared clusters remained largely unchanged. Selected clusters were enriched in TBLN only in the pH1N1 repertoire, but not in the PRCV Blg20. A possible explanation might be that the pH1N1 infection was more restricted to the lung, while PRCV Blg20 induced a generalised systemic immune response.

**Figure 3:**
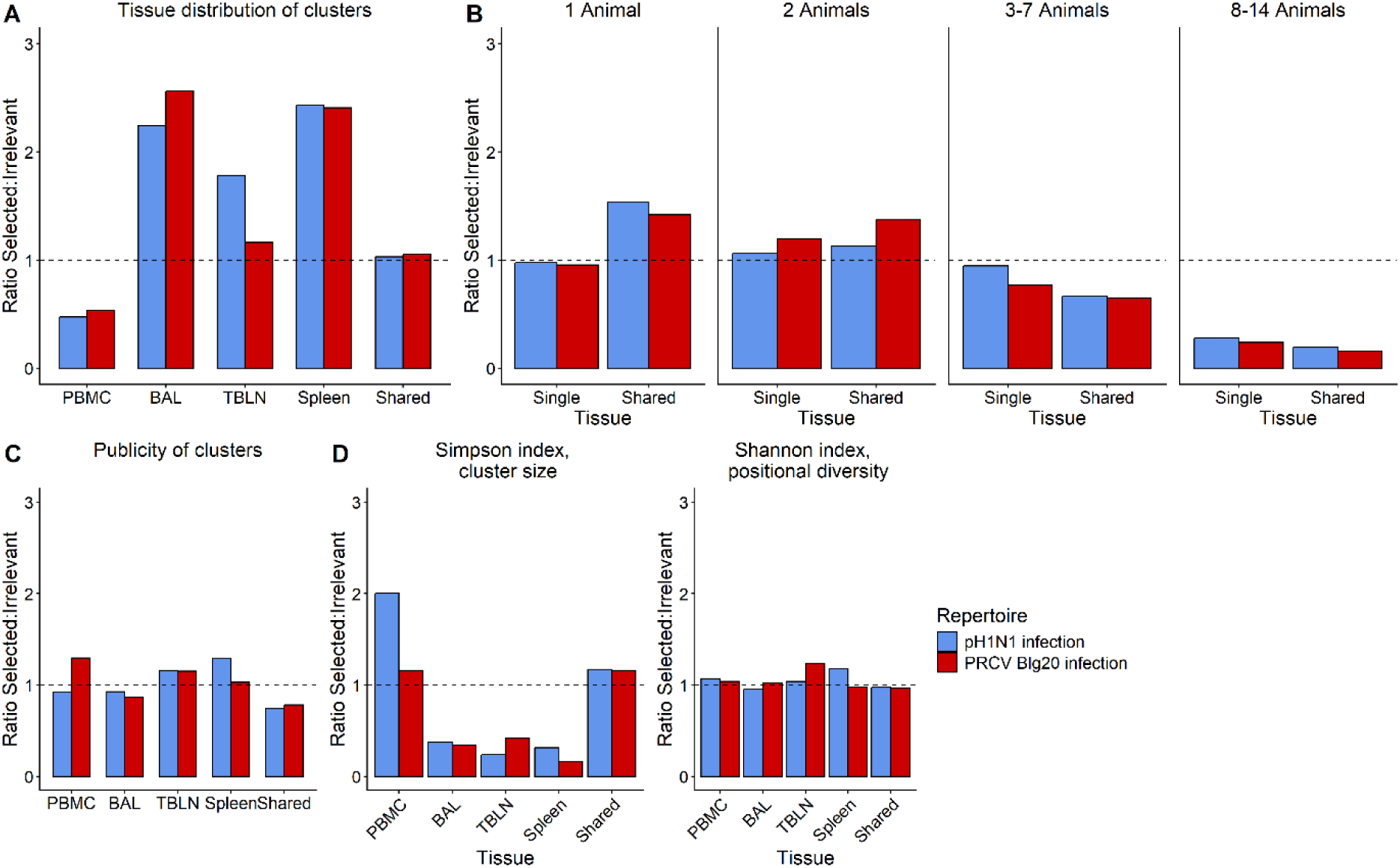
Comparison of antibody repertoires following pH1N1 or PRCV Blg20 infection Ratios between selected and irrelevant clusters were compared across the antibody repertoires following infection with either pH1N1 or PRCV Blg20. **A:** Ratio of the proportion of selected to irrelevant clusters restricted to a single tissue or detected across more than one tissue. **B:** Ratio of the proportion of public selected to public irrelevant clusters across tissue categories. **C:** Ratio of the proportion of selected to irrelevant clusters shared between two or more animals across tissue categories following infection with either pH1N1 or PRCV Blg20. **D:** Ratio of diversity between selected and irrelevant clusters, shown for cluster size (Simpson index) and centroid amino acid sequence diversity (Shannon index). In all panels, bars above the dotted line indicate enrichment in selected clusters, whilst bars below the dotted line indicate enrichment in irrelevant clusters.

Overall publicity patterns were similar between the two infections (**Fig. 3B**). In both datasets, selected clusters shared across tissues were more often private than irrelevant clusters. Selected clusters were more frequently shared between two animals (indicated by ratios of selected to irrelevant clusters above the dashed line), but less frequently shared across larger numbers of animals. In both infections, selected clusters in BAL and shared clusters were less public than irrelevant clusters, whereas selected clusters in TBLN were more often public in both datasets (**Fig. 3C**). However, patterns of publicity also showed distinct features. In the pH1N1 repertoire, selected PBMC clusters were less public than irrelevant clusters, whereas the opposite pattern was observed following PRCV Blg20 infection. In the spleen, selected clusters were more public in pH1N1 but no different to irrelevant clusters in PRCV Blg20. This indicates that, despite common tissue-associated patterns of publicity, inter-animal sharing in blood and spleen varied according to pathogen.

Patterns of diversity in cluster size and centroid amino acid sequence were also similar between pH1N1 and PRCV Blg20 infections (**Fig. 3D**). Selected clusters in BAL, TBLN and spleen were more diverse in size than the corresponding irrelevant clusters, indicated by ratio of selected vs irrelevant clusters less than 1. However, in the pH1N1 repertoire, PBMC-only clusters were less diverse than irrelevant clusters, indicated by an approximate 2-fold difference in Simpson index which was not observed in the PRCV Blg20 repertoire. In both repertoires, selected clusters were generally larger than irrelevant clusters (**Figs. 1E, 2E**). Positional diversity of the centroid amino acid sequence was increased in the selected clusters in both repertoires. However, the magnitude and location of these differences varied: In the pH1N1 dataset, increased diversity was most evident in spleen restricted clusters, whereas in the PRCV Blg20 dataset this was more pronounced in the TBLN.

V gene segment usage in both infections was dominated by IGHV1-4 and IGHV1-S2, which accounted for more than 50% of the repertoire, followed by IGHV1-15, IGHV1-8, and IGHV1-10. This pattern is consistent with previous reports, and no clear differences were observed between tissues or treatments (**Figs. 1F, 2F**)^1, 19-21^.

In summary, selected clusters were enriched in BAL and spleen and reduced in PBMC in both infections, indicating localisation of the response to lymphoid and site-specific compartments. Selected clusters were generally less public than irrelevant clusters, particularly when detected in multiple tissues, demonstrating that infection-associated responses were individual-specific. Diversity analyses further supported tissue-specific patterns of response. Despite some differences between pH1N1 and PRCV Blg20, including no TBLN enrichment in PRCV Blg20, the overall architecture of the response was similar between the two infections.

### Antibody repertoire following prime-boost immunisation against PRCV

Following analysis of infection-induced responses, antibody sequences were assessed in five animals immunised twice 4 weeks apart with an adenoviral vector encoding PRCV 135 spike (S) and nucleocapsid (N) and decorated with spike receptor-binding domain (RBD) Ad-(S-N)-RBD^22^ (**Supp. Fig. S2A**). PBMCs were collected at 0, 7, 14, 21, 25, 29, 36, and 39 days post-prime immunisation. Spike-binding B cells were isolated by FACS from PBMC samples isolated 4 and 11 days post-booster immunisation and their paired heavy and light chains sequenced for comparison with the bulk repertoires and generation of spike-specific monoclonal antibodies. Antibody repertoire was assessed from IgG and IgA heavy chain sequences at all timepoints. Following quality control of sequencing data (**Supp. Fig. S2B, C**), clusters from PBMC obtained post-immunisation were defined as selected where less than 10% of reads were derived from pre-bleed samples, and two peaks were observed: one between 7 and 21 days after the prime dose, and a second four days after the boost immunisation (**Supp. Fig. S2D**)^23^. All other clusters were defined as irrelevant (**Supp. Fig. S2E**). Clusters were analysed for isotype composition, publicity, and diversity.

Irrelevant clusters primarily contained sequences of a single isotype, either IgG or IgA (**Fig. 4A**). Selected clusters were also often restricted to one isotype, but more frequently contained both IgG and IgA sequences, with 1.7-fold more selected clusters than irrelevant containing both isotypes (Χ^2^ = 212.4, df = 9, p < .0001, **Supp. Table S7**), indicating representation across multiple class-switched isotypes. A higher proportion of selected clusters were public, with 1.6-fold more selected clusters than irrelevant containing sequences from all five animals containing sequences from two animals (p < 0.0001), (**Fig. 4B, Supp. Table S8**). Increased publicity in selected clusters indicated that prime-boost immunisation elicited a more convergent repertoire across animals, as compared with infected pigs.

**Figure 4:**
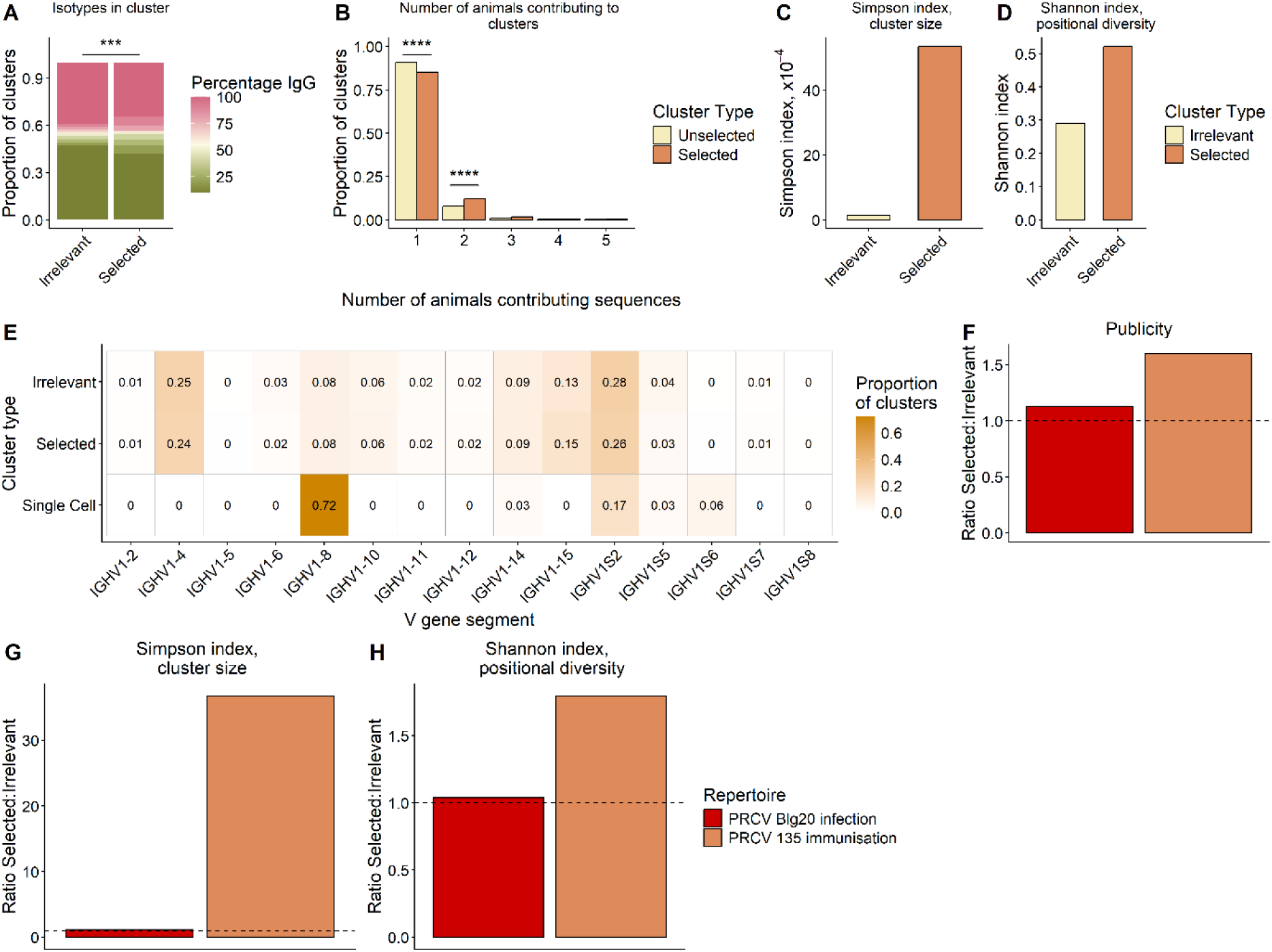
Antibody repertoire in PBMCs following prime-boost immunisation against PRCV 135 or infection with PRCV Blg20 Irrelevant and selected PBMC clusters were compared following PRCV 135 immunisation or PRCV Blg20 infection. **A:** Isotype usage in irrelevant and selected clusters. Isotype usage was analysed by Chi-square test (Χ2 = 212.4, df = 9, p < 2.2x10-16). **B:** Number of animals contributing sequences to each cluster. Distribution of sequences across 1-5 animals was compared by Fisher exact test for overall distribution and between cluster types for each number of animals. **C:** Cluster size diversity measured by Simpson index, where a higher value indicates lower diversity. **D:** Amino acid sequence diversity measured by Shannon index. **E:** V gene usage in centroid sequences of irrelevant and selected clusters following PRCV 135 immunisation, compared with antigen-binding B cells isolated by FACS (“Single Cell”). **F:** Ratio of PBMC-only clusters containing sequences from two or more animals. **G:** Ratio of cluster size diversity between selected and irrelevant PBMC-only clusters, measured by Simpson index. **H:** Ratio of amino acid sequence diversity between irrelevant and selected PBMC-only clusters, measured by Shannon index. In panels F-H, values greater than 1 indicate higher values in selected clusters than in irrelevant clusters. **** indicates p < 0.0001

Diversity in cluster size was decreased in selected compared to irrelevant clusters, indicated by increased Simpson index (**Fig. 4C**). In contrast, centroid amino acid sequence was more diverse in selected clusters as measured by Shannon index (**Fig. 4D**). Immunisation was therefore associated with diversification at the sequence level. Thus, although selected clusters following immunisation were more shared across animals and less diverse in size, they were more diverse in sequence composition, indicating that immunisation elicited similar overall responses in different animals while maintaining sequence diversity within those responses.

V gene segment usage was analysed in selected and irrelevant clusters following immunisation. As with infection the repertoire was dominated by IGHV1-4 and IGHV1-S2, followed by IGHV1-15, IGHV1-8, and IGHV1-10. In contrast to the bulk repertoire the single-cell heavy chains showed a strong bias towards IGHV1-8, followed by IGHV1-S2 (**Fig. 4E**). This enrichment of IGHV1-8 was not reflected in the selected clusters from the bulk repertoire, suggesting that bulk sequencing captures a broader and more heterogeneous pool of B cells, including non-antigen specific or low frequency clones. In contrast, single cell sorting enriches for high affinity antigen-specific populations, which may preferentially utilise particular V gene segments such as IGHV1-8 due to structural compatibility with dominant epitopes.

Together these data suggest that following prime-boost intramuscular immunisation, selected clusters in PBMC were more public and often contained both IgG and IgA, indicating a convergent, class switched response. These selected clusters showed reduced size diversity but increased sequence diversity, while V gene segment usage remained similar to infection, with enrichment of IGHV1-8 observed only in isolated antigen-specific B cells.

### Comparison of PRCV infection and immunisation repertoires

Repertoires following PRCV infection and immunisation were compared in clusters containing sequences from PBMC. In both infection and immunisation repertoires, a higher proportion of selected clusters than irrelevant clusters were shared between two or more animals (**Fig. 4F**). This increase was more pronounced following immunisation than infection (1.6-fold and 1.1-fold, respectively). Despite this, the overall proportion of public clusters was approximately 4-fold lower in the immunisation repertoire than the infection repertoire (p < 0.0001).

Selected clusters had lower size diversity than irrelevant clusters following immunisation, indicated by increased Simpson index (**Fig. 4G**). In contrast, diversity in amino acid sequence was 1.8-fold higher in selected clusters than irrelevant following immunisation but not infection (**Fig. 4H**).

### Isolation of PRCV spike-specific monoclonal antibodies

Isolation of mAbs enables direct assessment of antigen binding and neutralisation, complementing repertoire analysis with functional characterisation. We therefore isolated Spike binding IgG positive B cells by fluorescence activated cell sorting from BAL collected 12 days post-infection after PRCV Blg20 infection, and from PBMC collected 4 or 11 days after booster immunisation of Ad-(S-N)-RBD. Antibody variable regions were cloned and expressed as porcine IgG1. Only three mAbs were infection-derived whereas 66 were obtained following immunisation (**Supp. Fig. 3**). Of these, 45 immunisation-derived mAbs were selected for cloning (SP01- SP-45), together with all three infection-derived mAbs (SPi-01, SPi-02 and SPi-03).

Expression supernatants were screened by ELISA for binding to full length recombinant spike from PRCV Blg20 (PRCV Blg20 S) and ISU-1 (PRCV ISU-1 S) as well as the receptor binding domain from PRCV 135 (PRCV 135 RBD). Sequence diversity between the three recombinant protein is shown in **Supp. Fig. S4**. Of the 48 mAbs tested, 42 bound at least one spike protein, and 32 bound all three. A smaller subset showed restricted binding: two antibodies bound PRCV 135 RBD and PRCV Blg20 S, and four bound PRCV 135 RBD and PRCV ISU-1 S (**Fig. 5A**). Overall, 42 mAbs were spike-reactive, with many showing cross-reactivity between strains.

**Figure 5:**
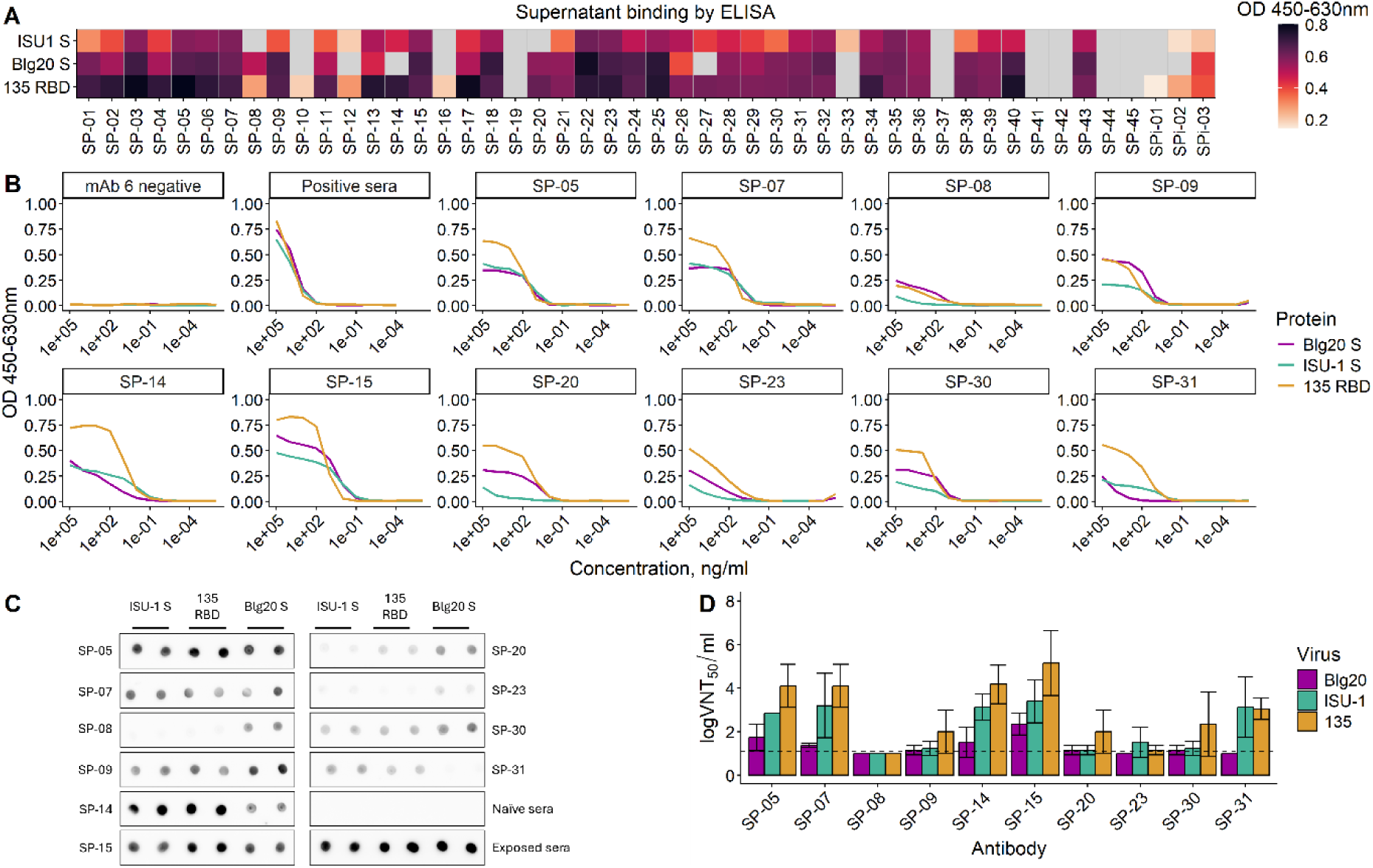
Characterisation of monoclonal antibodies isolated following PRCV infection or immunisation. Monoclonal antibodies were isolated from pigs immunised against PRCV 135 (SP-01 to SP-45) or infected with PRCV Blg20 (SPi-01 to SPi-03). **A:** Binding of antibody expression culture supernatants to recombinant full-length PRCV ISU-1 spike, full-length PRCV Blg20 spike and PRCV 135 receptor-binding domain (RBD), measured by ELISA. Grey squares indicate OD450 values within 3 standard deviations of the media-only control. **B:** Binding of ten purified antibodies to the three recombinant proteins measured by ELISA over a concentration range from 100 μg/ml to 0.1 pg/ml. Curves shown are representative of two independent experiments. **C:** Binding of purified antibodies to the three recombinant spike proteins assessed by dot blot at 1 μg/ml. **D:** Neutralising activity of purified antibodies against three PRCV strains, tested at concentrations up to 50 μg/ml. Bars at the dotted line indicate neutralisation below the limit of detection (VNT50/ml < 1.299).

Ten mAbs (SP-05, SP-07, SP-08, SP-09, SP-14, SP-15, SP-20, SP-23, SP-30 and SP-31) were selected for large-scale expression and purification to further characterise binding and neutralisation. These antibodies exhibited a range of binding, and each had distinct CDRH3 sequences. All purified mAbs bound PRCV135 RBD by ELISA but exhibited distinct binding profiles measured by IC50. SP-05, SP-07, and SP-15 had strong binding across all antigens, SP-09 and SP-30 had moderate binding, and SP-08 and SP-23 weak binding. In contrast, SP-14 and SP-31 displayed strong but strain-restricted binding, while SP-20 had limited and inconsistent binding (**Fig. 5B**)

Binding profiles by dot blot largely mirrored ELISA data, with some exceptions, likely reflecting differences in antigen conformation following immobilisation on plastic or nitrocellulose membrane (**Fig. 5C).** Neutralisation assays further distinguished functional activity. SP-05, SP-07, SP-14, SP-15, and SP-31 had strong neutralisation of PRCV 135, whereas SP-09, SP-20, SP-23 and SP-30 exhibited only minimal activity, and SP-08 showed none. Neutralisation of PRCV Blg20 was limited, with only SP-15 showing strong activity (**Fig. 5D).** The five antibodies exhibiting strong neutralisation activity against PRCV 135 also exhibited neutralisation activity against PRCV ISU-1, however activity against PRCV ISU-1 was consistently weaker than against PRCV 135.

Overall, based on their binding and neutralisation, the mAbs could be grouped into three categories (**Table 1**). Broad neutralisers (SP-05, SP-07, and SP-15) exhibited strong ELISA binding to all antigens, with corresponding dot blot profiles and robust neutralisation activity. In contrast, weak or non-neutralising mAbs (SP-08, SP-23, SP-09, and SP-30) had weak to moderate ELISA binding, largely consistent dot blot signals, and little to no detectable neutralisation. A third group comprised strain-restricted mAbs (SP-14, SP-31, and SP-20), which exhibited strong but antigen restricted ELISA binding, minor discrepancies in dot blot reactivity, and strain dependent or limited neutralisation activity. These results demonstrate substantial heterogeneity in both binding breadth and functional activity among mAbs targeting the RBD of PRCV 135.

Interestingly, none of the mAbs sequences mapped to the bulk heavy chain repertoire following either PRCV infection or immunisation. This suggests that the selected clustering approach captures only a subset of the antigen-associated response. However, this may also reflect the low abundance of bona fide antigen-specific clones in bulk datasets, their exclusion by read-count or kinetic thresholds, or enrichment of a distinct probe-binding subset by flow cytometric sorting.

## Discussion

This study demonstrates that respiratory viral infection and intramuscular immunisation generate distinct repertoire architectures in pigs, with infection producing localised and individualised responses and immunisation promoting greater convergence across animals. Following pH1N1 or PRCV Blg20 infection, response-associated selected clusters were consistently enriched in BAL and spleen, with reduced representation in peripheral blood. This supports a model in which B cell responses are primarily localised to sites of antigen exposure and secondary lymphoid organs, rather than being broadly reflected in the circulation. The similarity in tissue distribution between the two infections suggests that this spatial organisation is a general feature of respiratory immune responses, although the stronger enrichment in TBLN following pH1N1 indicates pathogen-specific differences in lymphoid involvement. This might be due to the more generalised response following PRCV Blg20 infection, while pH1N1 was mainly restricted to the respiratory tract^7, 24^. Our findings are consistent with observations from mouse influenza model, where respiratory infection with PR8 generated highly compartmentalised B cell responses, with distinct and largely non overlapping clones identified in lung, spleen, and lymph nodes supporting the concept that antigen-driven responses are shaped locally within tissues rather than uniformly distributed across the organism^25, 26^.

Publicity analysis further showed that infection-associated selected clusters were less shared across animals than irrelevant clusters. This indicates that recent antigen-driven responses are more individualised, likely reflecting differences in antigen exposure, germline repertoire, and stochastic aspects of clonal selection. The observation that irrelevant clusters were more public is consistent with the presence of common baseline clonotypes that are widely shared as naïve repertoires across individuals contain a core of shared or “public” sequences despite overall high diversity^27^. In contrast, antigen-experienced selected clusters were more private due to selection and expansion of distinct clones in each individual^28^. Despite some variation between tissues and pathogens, selected clusters were generally restricted to fewer animals, particularly when detected across multiple tissues, reinforcing the idea that functional responses are personalised.

In contrast to infection, prime boost intramuscular immunisation induced a more convergent PBMC repertoire across animals. Selected clusters were more frequently public, indicating that vaccination drives common responses, likely due to controlled and repeated antigen exposure. Such convergent or “public” antibody responses following vaccination has been reported, with shared clonotypes identified after influenza and SARS-CoV-2 immunisation^29-32^. At the same time, immunisation was associated with reduced cluster size diversity but increased sequence diversity, consistent with expansion of dominant clones together with ongoing diversification, a pattern also observed following repeated antigen exposure and boosting in vaccine settings^29^. The presence of both IgG and IgA within selected clusters further indicates that immunisation can engage multiple class switched compartments, in agreement with studies showing that vaccination can induce both systemic and mucosal class switched responses depending on antigen and regimen^33^. In our study, comparisons between PRCV infection and immunisation were limited to PBMC only, which represent a minority of the total B cell population. Future work should extend this analysis to different tissues, including after respiratory immunisation, to better define how route of antigen exposure shapes the repertoire across anatomical sites.

Studies of porcine antibody repertoires remain relatively limited, but several common features have emerged. In pigs, the immunoglobulin locus contains a relatively small number of functional V, D and J genes, meaning that repertoire diversification depends less on combinatorial diversity and more on junctional diversity and somatic hypermutation^15, 16, 34^. Early work therefore focused mainly on spectratyping and CDRH3 length distribution, which showed repertoire perturbations after infection with porcine reproductive and respiratory syndrome virus (PRRSV), swine influenza virus or porcine circovirus 2 (PCV2), but did not provide sequence-level resolution of the responding populations^17, 35^. More recent sequencing-based studies in porcine epidemic diarrhoea virus (PEDV) vaccination or infection also showed that porcine repertoires are dominated by a limited set of IGHV genes and that changes in bulk V-gene usage after antigen exposure may remain relatively modest^19, 20^. In the recent study by Herbet et al^30^, kinetic clustering of heavy-chain repertoires after pseudorabies virus vaccination and challenge identified numerous public clusters shared between pigs, indicating that convergent responses can emerge in this species despite restricted germline usage. Our findings are broadly consistent with this general framework, as we also observed limited changes in overall V-gene usage. However, our data further show that major differences in the porcine antibody response can be revealed through tissue distribution, inter-animal sharing and repertoire diversity. In this context, convergence appears to be one possible feature of porcine humoral immunity, but not a universal one, as the responses observed here following respiratory infection were more strongly characterised by tissue compartmentalisation and individuality, whereas immunisation promoted greater sharing across animals.

V gene segment usage showed no major changes across tissues or infection type and was dominated by a limited set of IGHV1 genes, including IGHV1-4 and IGHV1-S2, IGHV1-15, IGHV1-8, and IGHV1-10. This suggests that porcine antibody responses arise from a relatively fixed germline pool, with repertoire changes driven by clonal selection rather than gene recruitment, consistent with studies in PRRSV and PEDV infections and baseline analyses^1, 18, 19, 21^. In contrast, the enrichment of IGHV1-8 in Spike specific IgG positive B cells likely reflects antigen driven selection within a narrower functional compartment, not captured in bulk sequencing. This distinction between bulk and single cell data also aligns with reports in PEDV vaccination^20^, supporting overall stable germline usage with selective enrichment of specific V genes in antigen specific responses.

Repertoire sequencing provides a broad view of clonal expansion, diversity, and sharing, but does not directly define the functional properties of the antibodies involved. We therefore isolated mAbs to validate whether selected clusters correspond to antigen binding and neutralising activity, and to determine the relationship between repertoire and functional activity. Interestingly only three of the mAb sequences were detected within the bulk heavy chain repertoire after either PRCV infection or immunisation. This indicates that the clustering strategy identifies only a fraction of the antigen associated response and fails to capture all truly antigen specific clones. These findings indicate that cluster based bulk repertoire analysis and flow cytometric antigen specific single cell isolation are complementary rather than interchangeable approaches.

Few mAbs were recovered following infection, and none showed detectable neutralisation. In contrast, immunisation resulted in a larger number of spike-specific mAbs, including broadly neutralising mAbs. It is also possible that the limited number of B cell-derived mAbs recovered after infection reflects technical limitations, such as the quality or the low frequency of antigen-specific B cells in BAL samples. This possibility was not extensively evaluated across tissues and cannot be excluded. However, binding breadth did not predict neutralisation, indicating that epitope specificity and functional properties are not directly inferred from binding alone.

Monoclonal antibodies against the PEDV spike have been generated from porcine samples^36^, but most mAbs targeting porcine coronaviruses are produced in mice following exposure to recombinant protein^37, 38 39-41^. Mabs directed against the transmissible gastroenteritis virus (TGEV) spike typically bind to four antigenic sites, two of which (A and D) are retained in PRCV. Neutralising antibodies against TGEV often target antigenic site A^37^, suggesting that the neutralising antibodies identified here may bind the same region. However, further characterisation is required to define the specific epitopes bound by the mAbs described here.

Overall, these results show that respiratory infection with influenza virus or PRCV generates spatially compartmentalised, diverse, and highly individual-specific antibody repertoire responses, whereas intramuscular PRCV immunisation promotes greater convergence and shared responses across animals. Tissue localisation, particularly within the respiratory tract and draining lymph nodes, emerges as a key determinant of clonal expansion and diversity. Despite stable germline usage, functional antibody responses are shaped by local antigen exposure, clonal selection, and host specific factors, resulting in distinct repertoire architectures between infection and immunisation.

## Methods

### Cells, viruses and recombinant proteins

Influenza A/swine/Gent/53/2019 (pH1N1) and PRCV strain PRCV/swine/Belgium/PS-071/2020 (PRCV Blg20) were provided by K. Van Reeth (Ghent University). PRCV 135 strain 86/13508 was provided by S. Cartwright (Animal and Plant Health Agency, APHA)^42^, PRCV strain ISU-1 was purchased from BEI Resources (NR-43286). pH1N1 virus was cultured in MDCK cells^43^. PRCV viruses were cultured in ST cells using PRCV growth media (EMEM (Sigma-Aldrich) supplemented with 0.02% yeast extract, 10% tryptose-phosphate broth, 1% L-glutamine, 1% penicillin-streptomycin solution, and 1% nystatin solution). Viral titre was quantified by plaque assay in ST cells.

Swine testis (ST) cells were cultured in complete ST media (Advanced MEM (Gibco) supplemented with 5% FBS, 1% penicillin/streptomycin solution, and 1% L-glutamine). Cells were passaged once per week to no more than passage 30.

Expi293FTM cells were used for transient expression of antibodies. They were cultured in Expi293 Expression medium (Gibco, A1435101) to a density of 3-5x10^6^ viable cells/ml before passaging. Expi293FTM cells were cultured in a shaking incubator at 37°C, 5% CO_2_, and 120 rpm.

Three recombinant spike proteins were used to analyse the binding of the antibodies. The trimeric ectodomain of ISU-1 spike (PRCV ISU-1 S) was produced as described previously^14^. The trimeric ectodomain of PRCV Blg20 spike (PRCV Blg20 S) was expressed transiently in HEK-293F cells (CVCL-6642). To this end, human codon-optimized cDNA encoding the ectodomain of PRCV Blg20 S (GenScript) was cloned in frame with sequences encoding a GCN4-isoleucine-zipper trimerization motif^44^ a tandem repeat of a biotinylation acceptor peptide^45^ and a double Strep tag into a pCAGGS expression plasmid To stabilize the prefusion conformation of the spike protein two proline substitutions at positions 913 and 914 were introduced^46^ (Genbank PZ305746). The secreted S proteins were purified from cell culture supernatants by affinity chromatography using strep-tactin sepharose beads (IBA Lifesciences) followed by enzymatic biotinylation. The recombinant receptor binding domain from PRCV 135 (PRCV 135 RBD) included residues 300-449 of the PRCV 135 spike protein (accession number: URY50789.1), with the addition of an N-terminal IgK-leader sequence and a C-terminal C-tag. The protein was expressed in ExpiCHO-S cells and purified from supernatant by affinity chromatography using C-tag affinity resin, before dialysis into tris-buffered saline^22^.

### In vivo infection and immunisation studies in pigs

Samples from two *in vivo* studies were used for this work (**Supp. Figs. S1A**, **S2A**)^22^. All *in vivo* studies were performed in accordance with the U.K. Government Animal (Scientific Procedures) Act 1986 under Project License PP7764821and PP2064443 and were approved by ethical review processes at The Pirbright Institute and Vetquest. The Pirbright Institute conforms to Animal Research: Reporting of Animal Experiments guidelines. All animals were assessed for clinical signs including coughing, sneezing, nasal and eye discharge, rectal temperature, faecal consistency, appetite and demeanour. Endpoints based on pyrexia, behaviour, anorexia, and digestive or respiratory signs were specified in project licences PP7764821 and PP2064443. Virus was administered intranasally (1 ml per nostril) following sedation with 3 mg/kg Zoletil and 1.5 mg/kg Stresnil, and inoculum was backtitrated to confirm delivery of equivalent viral load to all animals.

Fifteen pigs were intranasally infected with 8x10^6^ plaque-forming units (PFU) each of either pH1N1 or PRCV Blg20. At 1, 5, and 12 days post-infection, 5 pigs per infection group and two uninfected pigs were humanely culled. Samples from pigs sacrificed at 5 and 12 days post-infection were used for antibody repertoire analysis. Peripheral blood mononuclear cells (PBMC), bronchoalveolar lavage (BAL), tracheobronchial lymph node (TBLN) and spleen samples were collected from all animals at postmortem, with an additional PBMC sample collected from the day 12 group at 7 days post-infection (**Supp. Fig. S1A**).

In the PRCV immunisation study, five pigs were immunised with replication-deficient adenovirus encoding full-length spike and nucleocapsid proteins and decorated with receptor binding domain from PRCV 135 using the DogTag/DogCatcher system, Ad-(S-N)-RBD135^22^. All animals received a homologous booster dose at 25 days post-first dose. PBMC samples were collected at the indicated times throughout the study (0, 7, 14, 21, 25, 29, 36 and 39 days post-prime immunisation) and used for antibody repertoire analysis (**Supp. Fig. 2A**).

### Library preparation for antibody repertoire analysis

RNA was extracted using the Qiagen RNEasy kit (Qiagen, 74104) from up to 1x10^7^ cryopreserved cells (from the infection study) or cells preserved in RNALater (for the immunisation study) as outlined in **Supp. Figure S1, S2**. RNA was isolated according to the manufacturer’s instructions, quantified by NanoDrop, and stored at -80°C. As the infection studies were performed at SAPO4 containment, Triton X-100 was added to buffer RLT to final concentration of 1%, to facilitate movement of samples out of virus-handling laboratories.

cDNA was synthesised from 50ng of RNA using SensiScript RT (Qiagen, 205213) supplemented with 1 μl per sample Random Primer 9 and Oligo dT(23)VN (New England Biolabs, S1254S and S1327S) and 0.25 μl per sample RNaseOUT (Invitrogen, 10777019) as described previously^47^. Samples were incubated with cDNA synthesis mix for an hour at 37°C and stored at -20°C until required.

Nested PCR was performed to amplify the IgG and IgA variable domain sequences using an adapted protocol from Illumina 16S sequencing. Primers (**Table 2**) were designed to amplify the immunoglobulin heavy chain variable region and a portion of the constant region (**Table 3, 4**). Overhangs facilitating addition of indexing primers were added in a second PCR (**Table 5, 6**), and Illumina indexing primers uniquely identifying each sample in a third PCR, as previously described^47^. Before and after the third PCR, samples were cleaned using AmpureXP beads (Beckman Coulter, A63880) at a sample:beads ratio of 1:0.65, according to manufacturer’s instructions. Presence of PCR product was confirmed before each bead clean step by agarose gel electrophoresis.

### Sequencing

Samples were sequenced as described previously^12, 47^. Briefly, samples were analysed using a TapeStation 4200 system and D1000 DNA ScreenTape (Agilent Technologies Inc). Samples were pooled and quantified by NEBNext Library Quantification Kit for Illumina (New England Biolabs) and Qubit fluorometer (Thermo Fisher Scientific). Libraries were sequenced on an Illumina NextSeq 2000, with a 40% PhiX control incorporated to increase sequence diversity. Initial quality control was performed using FastQC v0.12.1, then filtering and adapter trimming performed using Trim Galore! v0.6.10 as described previously. Paired-end reads were merged using FLASH v1.2.1. Sequences were annotated for gene usage, isotype and framework using IgBLASTn and IgMAT^48^. Sequences were clustered based on the entire variable region with a 94% identity threshold using UCLUST. Clustering was performed separately for each infection condition, with samples from the uninfected animals included in both analyses. Reads were then normalised to account for animal, time point and tissue sequencing depth variation. Cluster abundances were normalised over the total number of sequences per sample and expressed as percentage of reads.

### Analysis of antibody repertoire

Clusters were partitioned into irrelevant (those unlikely to be antigen-specific) and selected (those which may be antigen-specific) based on kinetics profiles^23^. Due to uncertainty in partitioning, clusters containing less than 10 reads were excluded from analysis.

In the pH1N1 and PRCV Blg20 infection repertoires, clusters containing at least 10% of reads from samples obtained prior to infection or from uninfected animals were defined as irrelevant. Clusters containing less than 10% of reads from samples obtained prior to infection or uninfected animals were defined as selected.

In the PRCV immunisation repertoire, selected clusters were defined as those for which less than 10% of reads originated from samples collected prior to prime immunisation with a peak in abundance between 7 and 21 days post-prime immunisation, and a second peak 29 days post-prime immunisation (4 days post-boost immunisation). Peak timepoints were selected based on previously described kinetics^23^.

### Analysis of cluster size and positional diversity

Diversity in cluster size and centroid amino acid sequence were quantified by Shannon and Simpson diversity, two approaches widely used in immune repertoire analysis to capture evenness and richness of clonotypes and sequence variability^49, 50^. Diversity was estimated after randomly subsampling sequences to a common depth across groups to control for sampling bias.

In the pH1N1 and PRCV Blg20 repertoires, indices were calculated for selected and irrelevant clusters containing sequences from single tissues only (PBMC, BAL, TBLN, or spleen) or containing sequences from two or more tissues (shared). For comparison to the immunisation repertoire, diversity indices were also calculated for all selected and irrelevant clusters containing sequences from PBMC in the PRCV Blg20 repertoire. In the immunisation repertoire, diversity indices were calculated for selected and irrelevant clusters.

Shannon entropy was calculated as 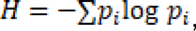 where 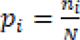 represents the proportion of sequences (cluster size) or amino acid frequency (positional diversity) in cluster *i* ^51^. Simpson diversity was computed as 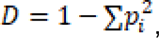 reflecting the probability that two randomly selected sequences belong to two different clusters^52^. Sequence alignment for positional diversity was performed using Clustal Omega^53^.

### Statistical analysis

Tissue distribution, isotype composition, and publicity of clusters was compared between selected and irrelevant clusters using Chi-square test with Yates’ continuity correction. Where expected values were less than 5, Fisher exact test was used. All statistical analysis was performed in R v4.5.2.

### Isolation of spike-specific B cells following PRCV infection and immunisation

To isolate spike-specific B cells, cryopreserved cells were stained for flow cytometry and sorted by fluorescence-activated cell sorting (FACS). Flow cytometry steps for screening experiments and controls for FACS sorts were performed in wells of U-bottom 96-well plates, whilst samples for sorting were stained in 15 ml Falcon tubes. Between each staining step, cells were washed twice with 200 μl per well or 1 ml per tube PBS containing 0.5% bovine serum albumin and 1 mM EDTA.

Cells were first stained with LIVE/DEAD^TM^ Fixable Near-IR Dead Cell Stain kit (Invitrogen, L34975), anti-porcine IgG-AF647 (Abbexa, abx142818), anti-porcine CD3ε-FITC (clone BB23-8E6-8C8, BD Bioscience, 559582), anti-porcine CD8α-FITC (clone 76-2-11, BD Bioscience, 551303), and anti-porcine CD172a-FITC (clone 74-22-15A, BD Bioscience, 561498). Antibodies in the first staining step were diluted in PBS and incubated with cells for 20 minutes at 4°C. Cells were then incubated with biotinylated Blg20 S diluted to 10μg/ml in wash buffer for 40 minutes at room temperature, followed by streptavidin conjugated to PE (EBiosciences, 12-431787) or BV421 (Biolegend, 405225) at 4°C for 30 minutes (**Table 7**). Antigen-binding B cells were defined as viable CD3ε^-^CD8α^-^CD172a^-^IgG^+^Spike-PE^+^Spike-BV421^+^ cells (**Supp. Fig. S3B**). Gating for spike protein was performed based on an uninfected, unimmunised control sample. Flow cytometry data was acquired using a BD LSR Fortessa, and sorting performed on a BD FACS Aria III Cell Sorter. Flow cytometry analysis was performed using FlowJo v10 (BD Bioscience).

To isolate spike-binding B cells post-infection with PRCV Blg20, a BAL sample isolated 12 days post-infection was selected based on previous flow cytometry experiments. 4x10^7^ cryopreserved cells were stained and single cells sorted into Hard-Shell 96-Well PCR plates (BioRad HSP9601) containing 10μl RNA isolation buffer (nuclease-free water containing 10mM Tris pH 8.0 and 1000 units/ml RNasin Plus Ribonuclease Inhibitor (Promega, N2611).

Screening was performed to identify samples for sorting post-immunisation against PRCV 135. Cryopreserved cells from samples isolated 25-39 days post-immunisation against PRCV 135 were stained as described above, and proportion of spike-binding cells determined. A sample from 29 days post-prime immunisation and a sample from 36 days post-prime immunisation were selected. Sorting of these samples was performed as for the post-infection sample.

During sorting experiments, sample and sorted cells were stored on ice until required for sorting or downstream analysis. cDNA synthesis was performed immediately wherever possible, or stored at -80°C.

### PCR amplification of IgG heavy and lambda or kappa light chains from single cells

cDNA synthesis was performed as for the antibody repertoire libraries. The IgG heavy chain, lambda light chain, and kappa light chain were amplified in nested PCRs as described previously^12^. Briefly, an initial PCR was performed to amplify the IgG heavy, lambda and kappa chains (**Table 4, 5**). A second PCR was performed to facilitate addition of unique indexes for sequencing (**Table 6, 7**), and presence of samples determined by agarose gel electrophoresis. Samples were tracked in a database, and wells containing an IgG heavy chain and a single light chain were selected for sequencing. Bead cleaning and indexing and sequencing was performed as for the antibody repertoire libraries^12^.

Heavy chain sequences were clustered with the antibody repertoire datasets. One sequence clustered with the post-immunisation antibody repertoire so this was selected for cloning. In addition, some sequences which did not cluster with the repertoire were selected.

### Expression and cloning of spike-specific monoclonal antibodies

All antibodies were expressed as porcine IgG1. Synthetic gene fragments encoding variable regions (Twist Bioscience) were inserted into pNeoSec_SS_VL_kappa, pNeoSec_SS_VL_lambda, or pNeoSec_SSFc_IgG1 vectors by InFusion cloning. Antibodies were transiently expressed in Expi293F^TM^ cells as described previously^12^.

At small scale, 0.5 μg each of heavy and light chain plasmid were incubated at room temperature for 10 minutes in 0.5 ml of Expi293 medium containing 0.5% polyethylenimine, before addition to 1x10^7^ Expi293F^TM^ cells in 5 ml of culture media. Valproic acid, sodium propionate and glucose were added 16 hours post-transfection to final concentrations of 0.76 mg/ml, 0.61 mg/ml, and 0.75% respectively. Cell supernatants were harvested 48 hours after addition of transfection additives by centrifugation at 2000 xg for 10 minutes.

A subset of antibodies was re-expressed at larger scale for more in-depth characterisation. Expression followed a similar protocol, with 50 μg of each plasmid added to 100 ml of cell culture. Antibodies expressed at larger scale were purified using a 1 ml HiTrap Protein G HP column and Äkta Pure chromatography system (Cytiva). Antibodies were eluted with 0.1 M glycine-HCl, which was neutralised immediately by addition of one tenth volume of 1M Tris-HCl. Exchange of buffer to PBS was achieved by dialysis overnight at 4°C using 10 kDa molecular weight cutoff cassettes (Thermo Fisher Scientific).

### Western blot

Presence of antibody in supernatant was confirmed by western blot. Cell culture supernatant was subjected to SDS-PAGE and transferred to nitrocellulose membranes using the iBlot2 system (Thermo Fisher Scientific). Membranes were blocked for 1 hour with PBS containing 0.05% Tween20 and 5% milk (western blocking buffer), then incubated for a further hour with anti-pig IgG-HRP (BioRad, AHP865P) diluted 1:20,000 in western blocking buffer. The membrane was washed three times with PBS containing 0.05% Tween20, then developed using ECL Prime reagent (Cytiva, RPN2232) and imaged using a GelDoc imager (Biorad).

### Enzyme-linked Immunosorbent Assay (ELISA)

ELISA was used to evaluate the culture supernatants or purified mAbs to recombinant Spike proteins. To assess binding, 96-well plates (half-area plates (Falcon) for culture supernatant or Maxisorp plates (Thermo Fisher Scientific) for purified antibody) were coated overnight at 4°C with 0.5 μg/ml recombinant ISU-1 S or PRCV Blg20 S, or 2 μg/ml 135 RBD diluted in PBS. Plates were washed three times with ELISA wash buffer (PBS containing 0.05% Tween20) then incubated for 1 hour at room temperature with ELISA block buffer (ELISA wash buffer containing 4% w/v Marvel milk powder). Samples were diluted 1:2 in ELISA block buffer (supernatants) or serially diluted 10-fold from 100 μg/ml (purified antibodies) and incubated on the plates for 1 hour at room temperature. Plates were washed three times with ELISA wash buffer, then incubated with goat anti-pig IgG-HRP (BioRad, AHP865P) diluted 1:20,000 in ELISA block buffer for 1 hour at room temperature. Plates were washed again, then developed with 25 μl per well (supernatants) or 50 μl per well (purified antibodies) of TMB substrate solution (Biolegend). Equivalent volume of 1 M sulfuric acid was added after four minutes (supernatants) or one minute (purified antibodies). Absorbance at 450 nm and 630 nm was detected using an ELx808 microplate reader.

### Dot blot

To assess binding of antibodies to recombinant proteins by dot blot, 100 ng recombinant full-length ISU-1 and PRCV Blg20 spike or 500 ng recombinant 135 RBD diluted in PBS was applied to a nitrocellulose membrane and dried until no visible moisture remained. Membranes were blocked using ELISA blocking buffer, then incubated with monoclonal antibody diluted to 1 μg/ml in ELISA blocking buffer. Membranes were washed 3 times for 5 minutes each with ELISA wash buffer, then incubated with goat anti-pig IgG-HRP (BioRad, AHP865P) diluted 1:5000 in ELISA block buffer. Membranes were washed a further 3 times for 5 minutes each, then developed using ECL prime reagent for 1 minute. All membranes were imaged simultaneously using a ChemiDoc imager (BioRad). All block steps and antibody incubations were performed for 1 hour at room temperature.

### Virus neutralisation assay

Neutralisation of antibodies against PRCV Blg20, PRCV 135 and PRCV ISU-1 was assessed in supernatant and purified mAbs by virus neutralisation assay. ST cells were cultured in 96-well plates to confluence. Culture supernatants were serially diluted 10-fold in PRCV growth media (EMEM supplemented with 0.02% yeast extract, 10% tryptose-phosphate broth, 2 mM L-glutamine, and 1% penicillin/streptomycin) starting at 1:5. Purified antibodies were serially diluted 5-fold in PRCV growth media starting at a concentration of 50 μg/ml. Serially diluted mAbs were incubated with 100 μl PRCV growth media containing 10^2^ plaque forming units of either PRCV Blg20, PRCV 135, or PRCV ISU-1 for 30 minutes at room temperature. After 30 minutes, ST cells were washed, the virus-antibody mixture applied, and cells incubated at 37°C for 48 hours. Cells were fixed with 4% formaldehyde for 1 hour at room temperature, then stained with 0.1% crystal violet for 20 minutes. Neutralisation titre resulting in 50% neutralisation per ml was calculated using the Reed-Muench endpoint calculation method^54^.

## Data availability

Antibody repertoire data is deposited in the European Nucleotide Archive.

## Acknowledgments

We are grateful to the animal staff at The Pirbright Institute for providing excellent animal care and the resources of the Bioinformatics Science Technology Platform. We thank Nancy Schuurman for technical support. This work was supported by the UKRI Biotechnology and Biological Sciences Research Council (BBSRC) IAA award BB/X019780/1 as part of the EPICVIR project at the international collaboration of research on infectious animal diseases (ICRAD) and Gates Foundation grant OPP1215550 (Pirbright Livestock Antibody Hub). Research at Pirbright was funded by BBSRC via the Pirbright Institute’s Strategic Programme Grants (ISPGs) [BBS/E/PI/230002A; BBS/E/PI/230002B and BBS/E/PI/230002C], BBSRC National Bioscience Research Infrastructure: High Containment and Low Containment Services and Science Platforms grants [BBS/E/PI/23NB0004, BBS/E/PI/23NB0003]. CR is supported by the Equal Opportunities Foundation (Hong Kong) and the Braithwaite Family Foundation. EJB was supported by a PhD studentship from the Pathogens and Host Defences DTP (University of Surrey and The Pirbright Institute). The funding sources had no involvement in study design, analysis, interpretation, or reporting of the results.

## Author contributions

**Conceptualisation:** Elma Tchilian

**Data curation:** Emily J. Briggs, Marie Bonnet-Di Placido, Valentine Chaillot, Bharti Mittal

**Formal analysis:** Emily J. Briggs, Marie Bonnet-Di Placido, Bharti Mittal

**Funding acquisition:** Elma Tchilian, Christine S. Rollier

**Investigation:** Emily J. Briggs, Shafiyeel Chowdhury, Bharti Mittal, Valentine Chaillot, Charlotte May, Kevan Hanson, Ehsan Sedaghat-Rostami, Sanis Wongborphid, Basudev Paudyal

**Methodology:** Emily J. Briggs, Bharti Mittal, Valentine Chaillot, Sarah Keep, Danish Munir, Marie Bonnet-Di Placido

**Resources:** Kevan Hanson, Cornelis A.M de Haan, Kristien Van Reeth, Matthew D. J. Dicks, Danish Munir

**Supervision:** Elma Tchilian, Christine S. Rollier, Marie Bonnet-Di Placido, Erica Bickerton, Sarah Keep

**Validation:** Emily J. Briggs, Shafiyeel Chowdhury, Bharti Mittal, Valentine Chaillot

**Writing – original draft:** Emily J. Briggs, Marie Bonnet-Di Placido, Elma Tchilian

**Writing – review & editing:** Emily J Briggs, Shafiyeel Chowdhury, Bharti Mittal, Valentine Chaillot, Charlotte May, Kevan Hanson, Ehsan Sedaghat-Rostami, Sanis Wongborphid, Erica Bickerton, Basudev Paudyal, Christine S. Rollier, Cornelis A.M. de Haan, Kristien Van Reeth, Sarah Keep, Matthew D. J. Dicks, Danish Munir, Marie Bonnet-Di Placido, Elma Tchilian

